# Chemogenetic Activation of Medial Prefrontal Cortex Projections to the Nucleus Accumbens Shell Suppresses Cocaine-Primed Reinstatement in EcoHIV Infected Mice

**DOI:** 10.1101/2024.12.20.629781

**Authors:** Qiaowei Xie, Mark D. Namba, Rohan Dasari, Lauren A. Buck, Christine M. Side, Samuel L. Goldberg, Kyewon Park, Joshua G. Jackson, Jacqueline M. Barker

## Abstract

HIV is highly comorbid with cocaine use disorder (CUD). Relapse is a major challenge in the treatment of CUD, and people living with HIV (PLWH) exhibit shorter time to relapse. One driver of relapse may be re-exposure to cocaine, which can be modeled in rodents using cocaine-primed reinstatement. This process involves neuroadaptations within the medial prefrontal cortex (mPFC) and nucleus accumbens (NAc) shell, regions that mediate cocaine reward learning and relapse-related behavior. HIV infection interacts with cocaine to alter corticostriatal circuits, which may further dysregulate cocaine seeking. To investigate the impact of HIV infection on cocaine reward learning and reinstatement and the role of mPFC-NAc circuits, we utilized the EcoHIV mouse model, a chimeric form of HIV-1 which can infect wild-type mice. Our findings demonstrate that EcoHIV infection enhances cocaine-primed reinstatement. We also observed increased cocaine-induced expression of the cellular activation marker cFos in the NAshell in EcoHIV-infected mice. Given the role of the mPFC-NAshell circuit in cocaine-seeking behaviors, we further demonstrated that chemogenetic activation of this circuit could reverse the behavioral deficits induced by EcoHIV. We propose that HIV infection contributes to neuroadaptations in the mPFC-NAshell circuit, and enhancing its activity may inhibit relapse-related behavior. These findings indicate that key neuronal circuits underlying cocaine reinstatement are similarly implicated in HIV infection and suggest potential strategies for managing relapse in PLWH.

## Introduction

Cocaine use disorder (CUD) and human immunodeficiency virus (HIV) infection are two global health crises that frequently co-occur (Dash et al., 2015; Lin et al., 2020). HIV infection and CUD possess independent but overlapping pathologies in the central neural system (CNS) and the immune system, which contribute to neurocognitive impairment and dysregulated reward seeking (Hauser et al., 2007; Wayman et al., 2016). Despite the high prevalence of co-occurring CUD and HIV infection, the consequences of HIV on cocaine seeking behaviors, and whether similar neural substrates regulate the expression of cocaine seeking in models of HIV infection, is not fully understood.

Relapse to cocaine seeking is a primary obstacle in treating CUD and a major factor contributing to cocaine overdose (Nnadi et al., 2005). During abstinence, factors including stress, drug-associated cues, or cocaine re-exposure can lead to relapse. The medial prefrontal cortex (mPFC) and nucleus accumbens (NAc) are essential regulators of craving and cocaine seeking (McFarland et al., 2003; Koya et al., 2009). Projections from the mPFC to NAc have distinct distributions and regulate different patterns of cocaine seeking behaviors: the prelimbic (PL) mPFC projection to NAcore is associated with reinstatement of drug seeking, while the infralimbic (IL) mPFC to NAshell projection inhibits reinstatement (McFarland et al., 2003; Peters et al., 2008).

Both cocaine and HIV infection disrupt neural substrates that mediate motivation, cognition, and reward processing. HIV infection is associated with corticostriatal circuit dysregulation that contributes to cognitive impairment in people living with HIV (PLWH) (Plessis et al., 2015; Wayman et al., 2016; McLaurin et al., 2018, 2021). This includes synaptic damage and neuronal loss in frontal cortex areas in HIV patients with HIV-associated dementia (Heaton et al., 1999; Ru and Tang, 2017; McLaurin et al., 2021). This is also observed in preclinical models of HIV infection. For example, HIV viral protein (e.g. gp120, Tat) exposure can directly impact N-methyl-D-aspartate receptor (NMDA) glutamate receptors by potentiating the phosphorylation and synaptic trafficking of NMDARs, leading to NMDA-mediated excitatory postsynaptic stimulation and increasing the synaptic damage by overactivation (O’Donnell et al., 2006; Ru and Tang, 2017). In addition, HIV-induced pro-inflammatory cytokine (e.g. TNF-α, IL-6 and IL-1β) release from infected/reactivated microglial cells/ macrophages can cause dysregulation of glutamatergic homeostasis and promote excitatory neuronal damage (New et al., 1998; Hauser et al., 2007; Namba et al., 2021). Thus, cocaine exposure in the context of HIV infection may accelerate impairment of the reward processing regions that lead to high relapse to cocaine seeking in PLWH.

Here, we investigate cocaine reward learning and reinstatement in a conditioned place preference (CPP) paradigm using the EcoHIV mouse model. In this model, the coding region of the HIV-1 glycoprotein gp120 is replaced with that of gp80, enabling infection of mice (Potash et al., 2005). Previous work demonstrated that EcoHIV-infected mice exhibit innate viral suppression (Gu et al., 2018), making this a potential model for studying the impact of HIV on reward-seeking behaviors in PLWH under viral suppression. Consistent with this, our previous findings identified neuroimmune alterations within the NAc that were not reversed by treatment antiretrovirals (Xie et al., 2024). Here, we find that EcoHIV infection did not impact cocaine CPP, but did impair extinction learning and increase cocaine-primed reinstatement. We also demonstrate that EcoHIV and cocaine interacted to alter cFos induction within the NAshell. We further demonstrate that chemogenetic activation of mPFC-NAshell projections prevented cocaine-primed reinstatement in EcoHIV-infected mice, similar to uninfected mice. Overall, the results from the current study indicate that EcoHIV infection altered cocaine-induced activity in the NAc and increased cocaine-primed reinstatement, and further, that corticostriatal pathways can be targeted to rescue cocaine relapse in populations with and without HIV infection.

## Materials and Methods

### Subjects

Adult male (n = 61) and female (n = 61) C57BL/6J mice (9 weeks of age upon arrival) were obtained from Jackson Laboratories. Following arrival, mice were group housed in same-sex cages for 7 days to acclimatize with *ad libitum* access to a standard chow diet and water. Mice were housed at the Drexel University College of Medicine under standard 12-hour light:12-hour dark conditions in microisolation conditions throughout the experiments. All experiments were approved by the Institutional Animal Care and Use Committee at Drexel University.

### EcoHIV-NDK production and inoculation

Plasmid DNA encoding the EcoHIV-NDK coding sequence (gift from Dr. David Volsky) was grown overnight in Stbl2 bacterial cells (ThermoFisher#10268019) and the plasmid DNA was purified using an endotoxin free plasmid purification kit (ZymoPure #D4200). Purified DNA was transfected into nearly confluent (80–90%)10 cm^2^ plates of low passage LentiX 293T cells (#632180, Takara Bio, San Jose, CA, USA), using a calcium phosphate transfection kit (Takara #631312). The cell culture supernatant was collected at 48 hours post-transfection using centrifugation at low speed (1500×g, 4°C), followed by passage through a cell strainer (40μm) to remove LentiX cells and cell debris. The supernatant, containing viral particles, was mixed 4:1 with a homemade lentiviral concentrator solution (4X; MD Anderson) composed of 40% (w/v) PEG-8000 and 1.2 M NaCl in PBS (pH 7.4). The supernatant–PEG mixture was incubated overnight at 4 °C on an orbital shaker (60 rpm). The mixture was centrifuged at 1500×g for 30 min at 4 °C. After centrifugation, the medium was removed, and the viral pellet was resuspended in cold, sterile PBS. The viral titer (p24 core antigen content) was determined initially using a LentiX GoStix Plus titration kit (#631280, Takara Bio, San Jose, CA, USA) and subsequently using an HIV p24 AlphaLISA detection kit (#AL291C, PerkinElmer, Waltman, MA, USA). Viral stocks were aliquoted and stored at −80 °C upon usage.

Following one week of acclimation, adult male and female mice were inoculated with 300 ng p24 equivalent EcoHIV-NDK or PBS (virus culture vehicle) as sham control by i.p. injection. To ensure housing consistency, all mice were singly housed. Blood samples were collected from all mice 1, 3, and 5 weeks following EcoHIV inoculation. Five weeks after inoculation, mice were assigned for behavioral training. This dose of virus and length of infection was selected as it produces systemic infection and immune response in the CNS and periphery (Potash et al., 2005; Kelschenbach et al., 2012, 2019; Gu et al., 2018; Namba et al., 2024; Xie et al., 2024). Following the completion of all experiments, to confirm the terminal EcoHIV-NDK infection status, spleens were isolated and flash-frozen on dry ice prior to perfusion and stored at -80C until processing. Splenic viral DNA burden was measured using Qiagen QIAamp DNA Mini Kit (#51304, Qiagen, Germantown, MD, US). Viral DNA was analyzed by the University of Pennsylvania Center for AIDS Research (CFAR). qPCR was conducted as described (Xie et al., 2024) using primers that amply sequences with HIV-LTR provided below. The OD value was detected by a NanoDrop™ spectrophotometer (Thermo Scientific) and used to determine the input cell numbers to normalize the data.

Kumar LTR F, GCCTCAATAAAGCTTGCCTTGA

Kumar LTR R, GGGCGCCACTGCTAGAGA

Kumar LTR Probe (FAM/BHQ), 5’CCAGAGTCACACAACAGACGGGCACA 3’

### Cocaine conditioned placed preference (CPP) and reinstatement test

Six weeks after inoculation with EcoHIV, mice were trained in a three-chamber CPP paradigm. Chambers had distinct walls (black, white, or gray) and floors (grid, wire mesh, or solid). The neutral gray chamber was in the middle of black and white conditioning chambers. Photocell beam breaks were used to calculate total time spent in each chamber, latency to enter, and locomotor behavior in the boxes using Med-PC V software (Xie et al., 2019).

Mice were habituated to the CPP room for at least 1 hour before the behavioral experiment. After habituation, mice were placed individually into the neutral chamber (middle gray chamber) with both door to black and white chamber opened (“Pre-test”). The ratio of time spent in the black and white chamber during the pretest was used to determine initial side bias, and the least preferred chamber was assigned as cocaine-paired chamber.

During conditioning, mice were assigned to receive cocaine (10mg/kg, i.p.) in the cocaine-paired chamber or saline injection in the opposite chamber on alternating days (**Figure 1A)**. Injections were administered prior to placement in the respective chambers, and mice were confined to the chamber for 30min in each conditioning session. This pattern of conditioning was performed for 4 consecutive days, resulting in a total of 4 pairings (2 cocaine and 2 saline). After conditioning sessions, mice were tested for the preference for the cocaine-paired chamber (“Post-test”). In this session, mice were placed in the neutral chamber with both doors retracted and allowed to freely explore all chambers for 20 minutes. The time spent in each chamber was assessed. CPP scores were calculated as [Post-test time spent in cocaine-paired chamber] – [Pre-test time spent in cocaine-paired chamber]. Following the CPP post-test, mice underwent 4 days of extinction training, which were identical to the CPP test sessions. Extinction scores were calculated as [Time spent in cocaine-paired chamber in each extinction sessions] – [Post-test time spent in cocaine-paired chamber].

**Figure 1.**
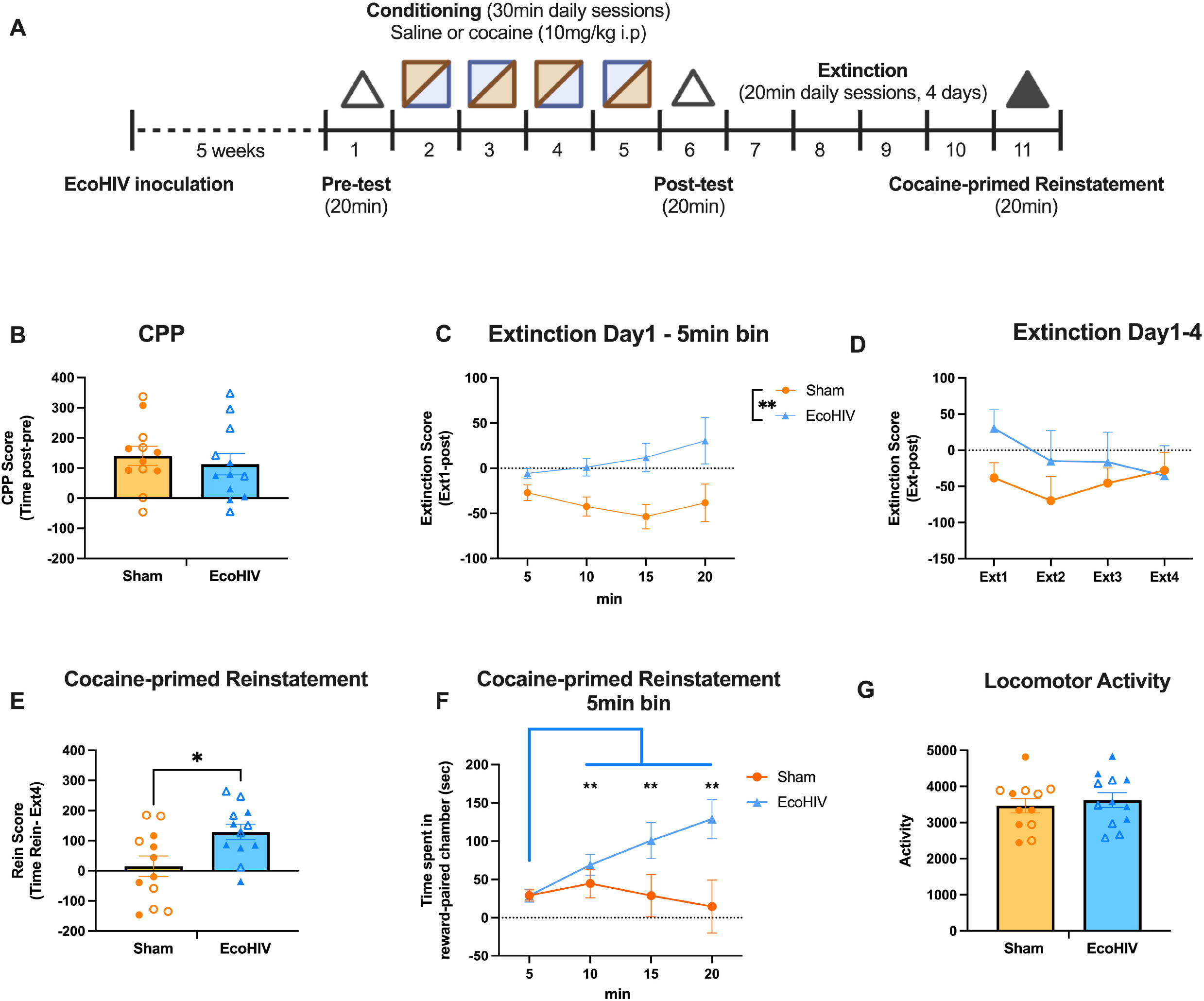
EcoHIV induced cocaine-primed reinstatement. (**A**) Timeline of experiment. Mice underwent EcoHIV or sham inoculation 5 weeks prior to training and testing in a cocaine CPP paradigm. Mice were tested for their CPP chamber bias during a 20-min pre-test session (open triangle). Cocaine (10 mg/kg, i.p.) and saline pairings (2 each, represented in alternating squares) occurred on alternating days followed by CPP the CPP post-test (open triangle). Mice then underwent 4 days of extinction training. In the cocaine-primed reinstatement test, mice received a cocaine injection (10mg/kg, i.p.) and were tested for 20 min (solid triangle). (**B**) A similar CPP was acquired in sham (n=12) and EcoHIV-infected mice (n=12). (**C**) To determine the effect of EcoHIV infection on within-session extinction acquisition, change in time spent in the cocaine-paired chamber was analyzed across 5-min bins on the first session of extinction. EcoHIV mice showed greater resistance to extinction (higher score) compared to sham mice. (**D**) To assess between-session acquisition of extinction, extinction scores were compared across days. EcoHIV and sham infected mice exhibited similar extinction across multiple training sessions. (**E**) EcoHIV-infected mice showed higher cocaine-primed reinstatement across the total session compared to sham mice. (**F**) EcoHIV, but not sham, mice exhibited increased reinstatement scores across the reinstatement test session. (**G**) No locomotor activity difference was observed between sham and EcoHIV-infected mice during cocaine-primed reinstatement. Timeline created with Biorender.com. Closed and open symbols represent male and female mice, respectively. Bars represent means ± SEM. ^*^p < 0.05, **p<0.01.

In the cocaine-primed reinstatement test, mice received a cocaine injection (10mg/kg, i.p.) and were tested for 20 min. The reinstatement scores were calculated as [Reinstatement time spent in cocaine-paired chamber] – [Extinction Day4 time spent in cocaine-paired chamber]. For the yohimbine-induced reinstatement test, mice underwent the same CPP and extinction training but received a yohimbine injection (2mg/kg, i.p.) followed by the 20 min reinstatement test.

### Immunohistochemical assessment of cFos

At least 48 hours after the cocaine reinstatement test, mice received an injection of cocaine (10mg/kg) and underwent transcardial perfusion with 4% paraformaldehyde (PFA) in 1X PBS 90 min later to assess cocaine-induced cFos expression in the mPFC and NAc. Fixed brains were collected and sectioned into 40 µm thick slices into triplicate wells for immunohistochemistry. Sections were blocked using 5% normal donkey serum (NDS) for 1 hour, then incubated in primary (rabbit anti-cFos, 1:10,000, Cell Signaling, 2250S) with 5% NDS overnight. Sections were then washed and incubated in donkey anti-rabbit secondary (1:1000, Jackson ImmunoResearch, 711-065-152) for 30 mins. The signal was amplified with an avidin-biotin complex (1:500, ABC kit, Vector Laboratories, PK-6200), then reacted with diaminobenzidine (Vector Laboratories, SK-4100) with nickel enhancement. Tissue was mounted and coverslipped with DPX (Electron Microscopy, 13512). Photomicrographs were obtained from three anatomical sections of the NAc (bregma: AP: +0.98mm, +1.21mm, +1.42mm) and mPFC (bregma: AP: +1.41mm, +1.69mm, +1.93mm) using a light microscope (Nikon, 10X). The images were imported into ImageJ for analysis. The average counts per area (total counts normalized to the selected area, measured as counts/mm^*2*^) across the three sections for both the NAc and mPFC were analyzed.

### Chemogenetic activation of mPFC-NAshell Gq signaling

A separate cohort of mice was used to determine the effect of chemogenetic activation of mPFC to NAshell projections on cocaine-primed reinstatement. Male and female were bilaterally injected into the mPFC (bregma: AP +1.8 mm, ML ±0.3 mm, DV –3.0 mm) with either pAAV-hSyn-hM3D(Gq)-mCherry (0.2 μL per side, 0.1μL/min, 5min diffusion, ordered from Addgene, 50474-AAV8), or control virus pAAV-hSyn-mCherry (114472-AAV8). Then a bilateral guide cannula was implanted into the NAshell (bregma: AP +1.4 mm, ML ±0.6 mm, DV –3.7mm). Mice were recovered for 7 days then underwent EcoHIV inoculation and cocaine CPP test as mentioned above. CNO infusion (500 μM, 0.2 μL/side, 0.1 μL/min) was performed over 2 min, followed by diffusion for 2 min. Mice were returned to the home cage for 10min before the cocaine-primed reinstatement test.

### Immunohistochemical confirmation of AAV placement

To verify AAV expression in the mPFC and canulation placement in the NAshell, 40 µm brain slices from control virus-or DREADD virus-expressing mice underwent RFP (for mCherry tag) immunofluorescent staining and were examined under an epifluorescence microscope. Slices were incubated in chicken anti-RFP primary (1:1,000, Novus Biologicals, Centennial, CO, NBP1-97371) overnight, then Alexa Fluor® 594 donkey anti-chicken secondary (1:250, Jackson ImmunoResearch, 703-585-155) for two hours. Mice with RFP signal in the mPFC region between 1.98mm and 1.34mm anterior to bregma and cannula placements in the NAshell between 0.98mm and 1.70mm anterior to bregma were included. Nine mice were excluded due to incorrect cannula placement in NAshell or insufficient DREADD expression in mPFC.

### Statistical Analyses

GraphPad Prism (10) was used for statistical analysis of behavioral and molecular data. Data were analyzed using unpaired *t*-test or two-way repeated measures (rm) ANOVA. Data with significant interactions were followed using Holm-Šídák’s multiple comparisons post hoc analysis, or for planned comparisons Fisher’s LSD multiple comparisons test where appropriate. The Greenhouse–Geisser correction was used if data violated sphericity. Statistical significance levels for each test were at p < 0.05. All data are displayed in figures as mean + standard error of the mean (SEM). Data will be available upon request.

## Results

### EcoHIV did not impact cocaine preference

To investigate whether EcoHIV infection impacted the development of cocaine CPP, we assessed the time spent in the cocaine-paired chamber after 4 conditioning sessions (2 cocaine and 2 saline) (**Figure 1A**). We found that following conditioning, both sham and EcoHIV-infected-mice demonstrated similar cocaine CPP [t(22)=0.5913, p = 0.5603; **Figure 1B**].

### EcoHIV-infected mice display impaired extinction learning

To investigate within-session extinction, change in time spent in cocaine-paired chamber was analyzed across 5-min bins on first session of extinction (**Figure 1C**). rm ANOVA revealed that EcoHIV mice had a higher extinction score compared to sham mice [main effect of EcoHIV: F (1, 22) = 8.211, p = 0.009], but there was no effect of time [F (1.261, 27.74) = 1.208, p = 0.2940, Greenhouse-Geisser corrected] or an interaction between EcoHIV and time [F (3, 66) = 2.305, p = 0.0848], suggesting that EcoHIV-infected mice exhibited persistent cocaine seeking across the Day 1 extinction session. There was no effect of EcoHIV on between-session extinction [Two-way rmANOVA, main effect of EcoHIV: F (1, 22) = 0.8234, p = 0.374; main effect of days: F (2.807, 61.76) = 1.521, p = 0.22, interaction: F (3, 66) = 1.597, p = 0.1985, Greenhouse-Geisser corrected; **Figure 1D**]. These data suggest that EcoHIV-infected mice exhibit delayed extinction learning compared to control mice but ultimately successfully extinguish cocaine seeking.

### EcoHIV infection significantly increased cocaine-primed reinstatement

Following extinction training, one cohort of mice underwent a cocaine-primed reinstatement test. An unpaired two-tailed t-test revealed that EcoHIV-infected mice exhibited greater cocaine-primed reinstatement scores than sham mice [t(22) = 2.649, p = 0.0147; **Figure 1E**], consistent with increased sensitivity to cocaine-primed reinstatement. To further examine this effect, we assessed the change in time spent in cocaine-paired chamber across 5-min bins which revealed that EcoHIV mice produced a higher reinstatement of cocaine seeking compared to sham mice (**Figure 1F**). A two-way rmANOVA revealed a significant interaction between time and EcoHIV [F (3, 66) = 8.231, p < 0.0001], as well as a main effect of time [F (1.310, 28.82) = 4.504, p= 0.0332, Greenhouse-Geisser corrected]. No main effect of EcoHIV was observed [F (1, 22) = 3.807, p = 0.0639]. Post hoc Holm-Šídák’s multiple comparisons revealed that EcoHIV-infected mice demonstrated increased preference scores at 10 (p = 0.0027), 15 (p = 0.0037) and 20-minutes compared to the start of the session (p = 0.0027). Sham mice did not exhibit an increase in preference score (all p’s > 0.6) Locomotor activity during the reinstatement test was assessed, but no differences in locomotor activity were observed between EcoHIV and sham mice [t(22) = 0.5371, p = 0.5966; **Figure 1G**], demonstrating increased reinstatement of cocaine seeking was not due to acute hyperlocomotor response-induced by cocaine in EcoHIV-infected mice.

A separate cohort of mice underwent yohimbine-induced reinstatement. While mice established a similar CPP (**Figure 2A**), in contrast to cocaine-primed reinstatement, neither sham nor EcoHIV-infected mice exhibited yohimbine-primed reinstatement [one sample t-tests vs 0, sham: t(11) = 1.041, p = 0.3204; EcoHIV: t(11) = 0.2523, p = 0.8055]. Further, EcoHIV infection did not impact yohimbine-induced reinstatement [unpaired t-test: t(22) = 0.7700, p = 0.4495, **Figure 2B**], suggesting that potentiating effects of EcoHIV infection may be selective to drug-primed, but not yohimbine-primed, reinstatement. A two-tailed, unpaired t-test revealed no differences in locomotor activity between sham- and EcoHIV infected mice during yohimbine-induced reinstatement [t(22) = 0.2188, p = 0.8288, **Figure 2C**].

**Figure 2.**
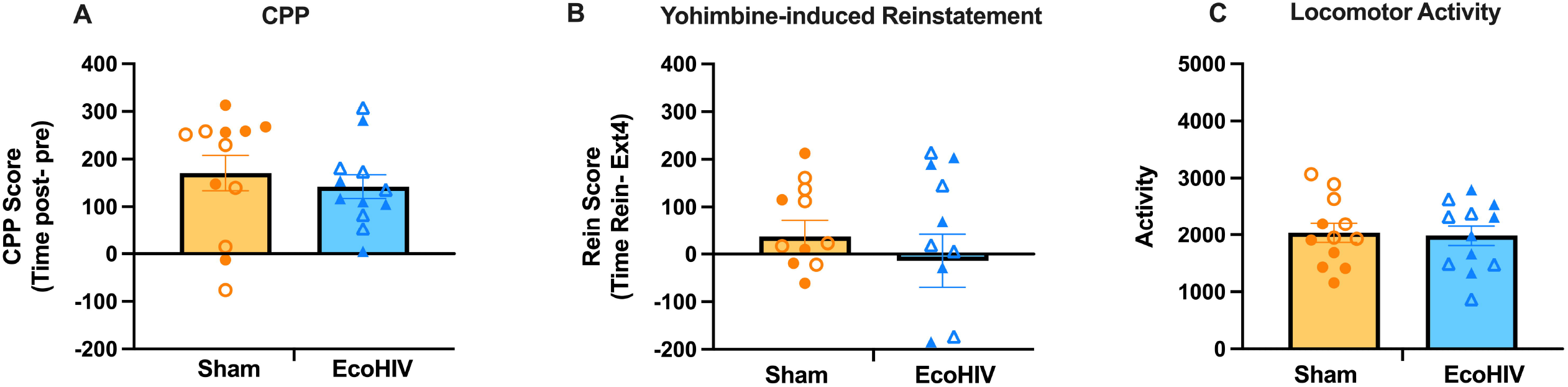
EcoHIV did not impact yohimbine-induced reinstatement. (**A**) A similar CPP was acquired in sham (n=12) and EcoHIV-infected mice (n=12). (**B**) No difference was observed between sham and EcoHIV-infected mice in yohimbine-induced reinstatement test. (**C**) No locomotor activity difference was observed between sham and EcoHIV-infected mice after yohimbine injection during reinstatement. Closed and open symbols represent male and female mice, respectively. Bars represent means ± SEM.

### EcoHIV infection increased cFos induction in the NAshell

To determine if cocaine had differential impacts on putative cellular activity within the mPFC and NAc of EcoHIV and sham mice, expression of the protein product of the cellular activity marker cFos was measured in mPFC and NAc subregions following an acute cocaine or saline injection. A two-way ANOVA indicated a main effect of cocaine [F (1, 19) = 7.265, p = 0.0143] and a main effect of EcoHIV [F (1, 19) = 5.094, p= 0.0360] in upregulating cFos expression in the NAshell. No interaction effect was observed [F (1, 19) = 2.973, p=0.1009; **Figure 3A**]. While cocaine also increased cFos expression in the NAcore [F (1, 19) = 4.772, p = 0.0417], there was no effect of EcoHIV [F (1, 19) = 0.02334, p = 0.8802] or interaction between cocaine and EcoHIV in the NAcore [F (1, 19) = 0.1771, p = 0.6786; **Figure 3B**]. No cFos expression differences were observed in IL [main effect of cocaine: F (1, 19) = 0.8650, p = 0.3640, main effect of EcoHIV: F (1, 19) = 0.007179, p = 0.9334, interaction: F (1, 19) = 0.04101, p = 0.8417; **Figure 3C**] or in PrL [main effect of cocaine: F (1, 19) = 1.035, p = 0.3219, main effect of EcoHIV: F (1, 19) = 0.8100, p = 0.3794, interaction: F (1, 19) = 0.1730, p = 0.6821; **Figure 3D**].

**Figure 3.**
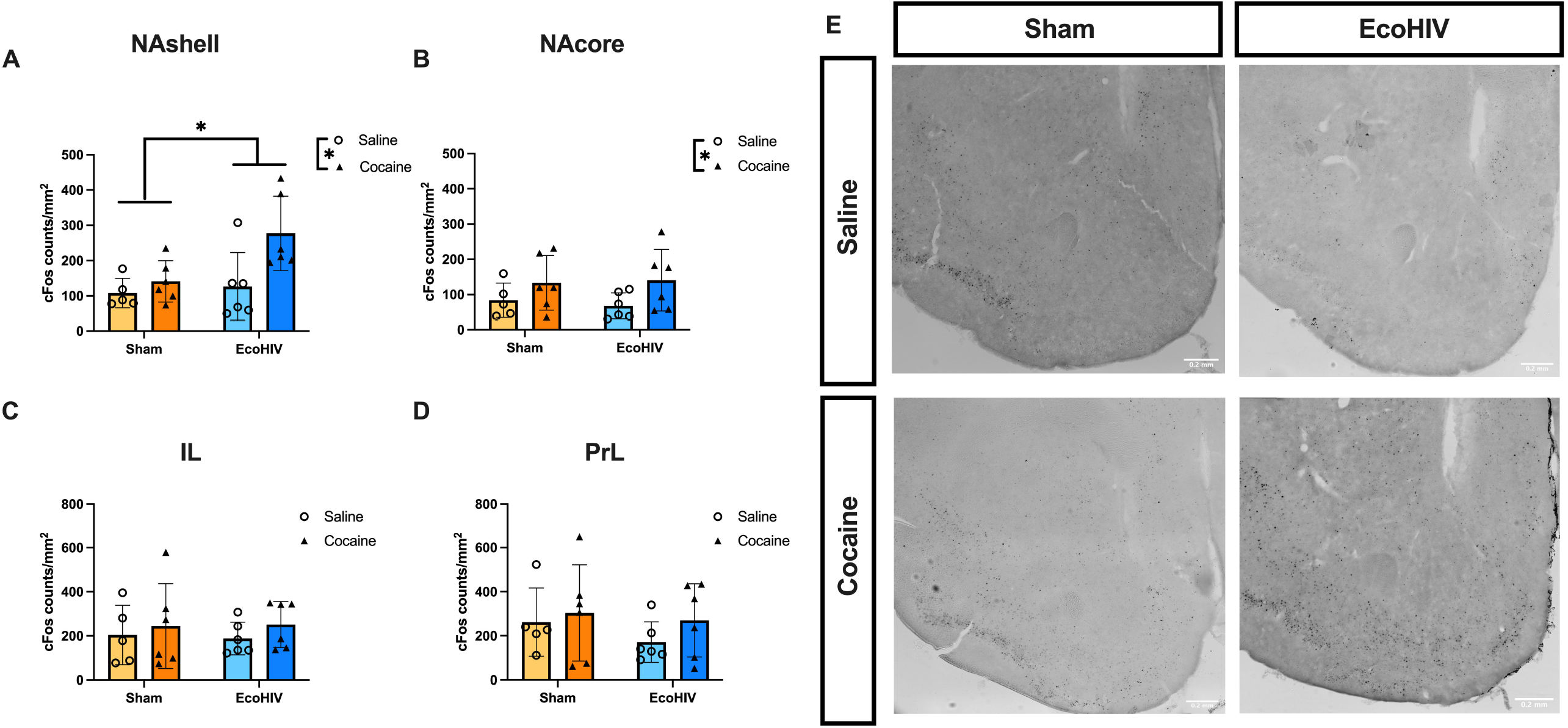
Expression of cFos in subregions of the mPFC and NAc. Cocaine and EcoHIV interacted to increase cFos expression (counts/mm^2^) in the NAshell (**A**), while cocaine increased cFos expression in the NAcore (**B**). No effect of EcoHIV or cocaine was observed in IL (**C**) or PrL (**D**). (**E**) Representative immunohistochemistry images of cFos in the NAc. n=6/group Bars represent mean +/-SEM. ^*^p<0.05

### Chemogenetic activation of Gq signaling within mPFC to NAshell circuits suppresses cocaine-primed reinstatement in EcoHIV-infected mice

As the mPFC projections to the NAshell are known to suppress reinstatement, we assessed the ability of chemogenetic activation of mPFC projections to the NAshell to suppress cocaine-primed reinstatement in EcoHIV infected mice (Timeline: **Figure 4A**). A two-way ANOVA revealed a significant main effect of DREADD [F(1, 43) = 4.300, p = 0.0441] on reinstatement scores, but no main effect of EcoHIV [F(1, 43) = 0.02411, p = 0.8773] or interaction between DREADD and EcoHIV [F(1, 43) = 1.713, p = 0.1976, **Figure 4B**], indicating DREADD activation suppressed reinstatement in both sham and EcoHIV infected mice. No difference was observed in locomotor activity during reinstatement between the 4 groups (**Figure 4C**), indicating the DREADD-suppressed reinstatement was not associated with locomotor suppression. These findings suggested chemogenetic activation of NAc-projecting mPFC neurons suppressed cocaine-primed reinstatement independent of EcoHIV infection status.

**Figure 4.**
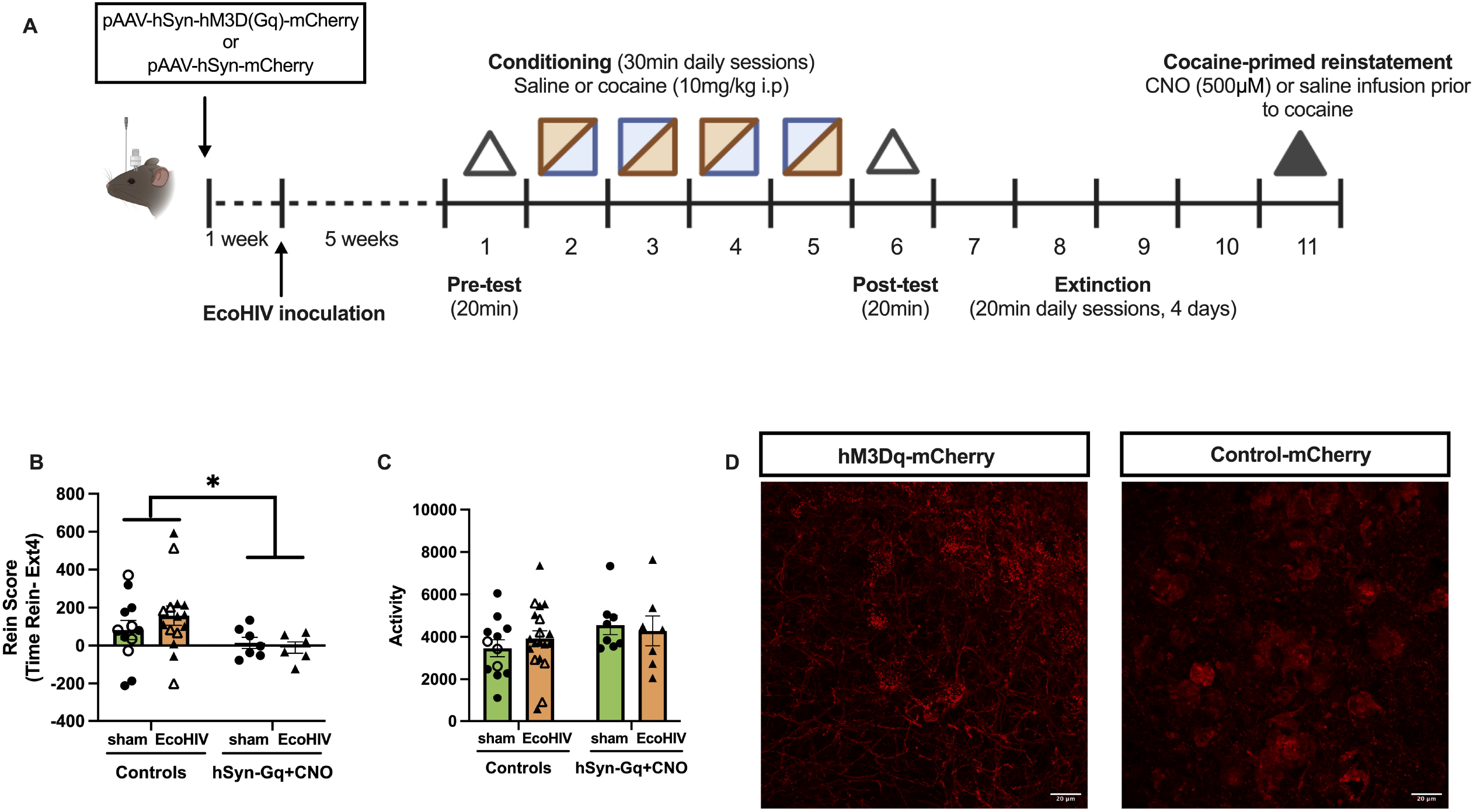
Chemogenetic activation of mPFC-NAshell projections. **(A)** Timeline of DREADD study. Anterograde neuronal Gq-DREADD (pAAV-hSyn-hM3D(Gq)-mCherry) or control (pAAV-hSyn-mCherry) virus was injected to the mPFC of mice prior to EcoHIV inoculation. Five weeks after inoculation, mice underwent the cocaine CPP and cocaine-primed reinstatement test. On the day of the reinstatement test, mice received an injection of CNO (or vehicle) to activate the mPFC-NAshell circuit prior to cocaine injection. (**B**) Gq DREADD activation by CNO administration significantly attenuated cocaine-primed reinstatement in both sham and EcoHIV-infected mice. (**C**) No difference was observed in the locomotor activity during the reinstatement across four groups. (**D**) Representative immunofluorescence images of hM3Dq-mCherry or mCherry in the mPFC sections. Data represent mean +/-SEM, Closed and open symbols represent CNO and saline treatment, respectively. n= 7-9 /group. ^*^p < 0.05.

### EcoHIV infection validation

To validate the infection status of mice, we isolated and purified DNA from the spleens of all EcoHIV-infected mice and a randomly selected subset of sham mice, and quantified HIV-1 long terminal repeat (LTR) DNA levels via qPCR. All EcoHIV-infected mice included in the dataset showed detectable levels of viral DNA. No viral DNA was detected in sham control mice. We verified that EcoHIV LTR DNA levels were in the range from 100 to 1500 viral DNA copies per 10^6^ spleen cells at the end of experiments (8 weeks following inoculation) (**Figure 5**).

**Figure 5.**
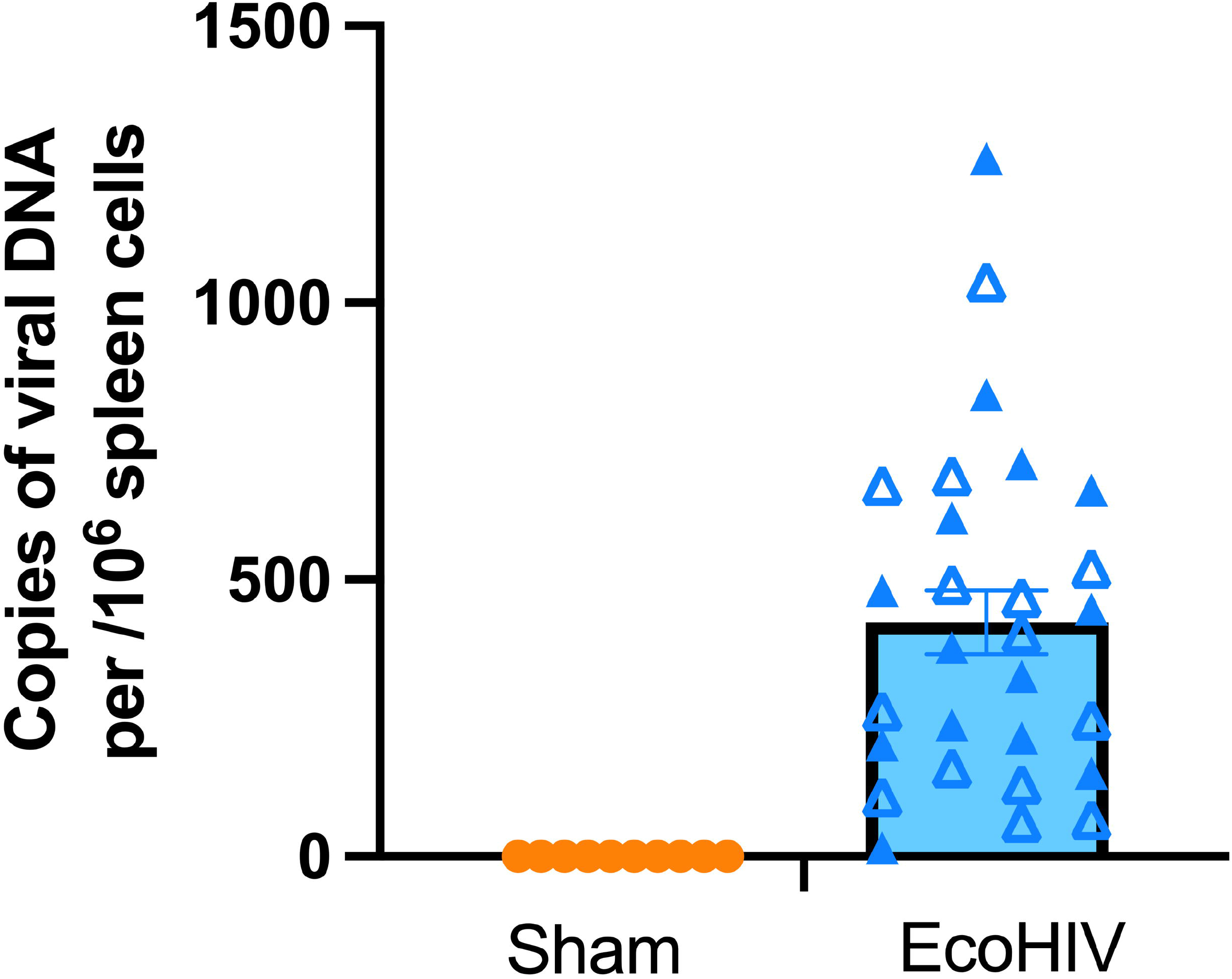
EcoHIV splenic DNA viral load. Copies of EcoHIV-NDK spleen viral DNA at the end of experiment (week 8 of infection). The number of copies were in the range from 100 to 1500 viral DNA copies per 10^6^ spleen cells. Closed and open symbols represent male and female mice, respectively. Data represent means ± SEM.

## Discussion

Cocaine use disorder is highly comorbid with HIV infection, and infection may alter the neural substrates regulating cocaine seeking. Our findings demonstrated that EcoHIV infection impaired cocaine extinction learning and potentiated cocaine-primed reinstatement but not yohimbine-induced reinstatement. Given that the mPFC-NAc circuit is sensitive to both cocaine and HIV, and plays a role in cocaine relapse after extinction (Peters et al., 2008; LaLumiere et al., 2010; Lalumiere et al., 2012; Meyer et al., 2014; Wayman et al., 2016), we investigated cellular activation in the mPFC and NAc subregions following acute cocaine exposure through cFos expression analysis. We found an interactive increase in cFos expression in the NAshell in EcoHIV-infected mice treated with cocaine, indicating greater sensitivity to cocaine treatment following infection. Moreover, chemogenetic activation of the mPFC-NAshell circuit reduced cocaine-primed reinstatement in EcoHIV-infected mice, suggesting that – as is observed in the absence of infection - activation of this pathway can suppress reinstatement.

Cocaine re-exposure is a common factor for relapse in people with CUD (Baum et al., 2009; Mimiaga et al., 2013). A priming dose of cocaine is often sufficient to trigger craving and relapse by reactivating extinguished memories formed through maladaptive plasticity during cocaine exposure (Farrell et al., 2018). The NAshell plays an important role in inhibiting cocaine-primed and cue-induced reinstatement, and its activity is required for extinction-induced suppression of cocaine seeking (Di Ciano et al., 2007; Peters et al., 2008). Evidence suggests that inactivation of the NAshell does not block cocaine reinstatement in the absence of extinction training, highlighting its specific role in extinction-driven inhibition of drug-seeking behavior (Peters et al., 2008; Lalumiere et al., 2012). Cocaine exposure and withdrawal has profound molecular and synaptic altercations in NAc. Our results show that EcoHIV-infected mice were resistant to initial acquisition of within-session extinction, spending more time in the cocaine-paired chamber compared to the post-test (**Figure 1C**). Additionally, EcoHIV-infected mice exhibited a higher rate of cocaine-primed reinstatement than sham controls. We observed increased cFos expression in the NAshell in response to cocaine in EcoHIV-infected mice, suggesting that EcoHIV infection may induce neuroplastic changes in these regions associated with behavioral alterations.

Glutamatergic plasticity has been linked to extinction learning. Extinction training induces an increase in AMPA glutamate receptor expression in the NAc, facilitating the retrieval of drug-associated memories and enhancing inhibitory control during relapse (Sutton et al., 2003; Self and Choi, 2004; Panopoulou and Schlüter, 2022). This upregulation of AMPA receptors may serve as a compensatory mechanism to counteract the decreased basal glutamate levels during cocaine withdrawal (Anderson et al., 2008; Kalivas, 2009). Repeated cocaine exposure with extinction training enhances AMPA and NMDA receptor trafficking to synaptic membranes and elevates glutamate transmission in the NAc, priming long-term potentiation (LTP) and promoting reinstatement of cocaine-seeking when re-exposed to the drug (Yao et al., 2004; Panopoulou and Schlüter, 2022). Indeed, HIV infection may exacerbate glutamatergic adaptations in the NAc. For example, studies show that HIV viral proteins promote NMDA receptor phosphorylation and potentiate glutamate excitotoxicity, leading to neuronal apoptosis (Haughey et al., 2001; Eugenin et al., 2007; Gorska and Eugenin, 2020). Previous work from our lab also reported that cocaine-seeking behavior is positively correlated with NMDA receptor (GluN2A) levels in the NAc, but this relationship is disrupted by EcoHIV infection (Namba et al., 2024). These findings suggest that EcoHIV may disrupt glutamatergic neuroadaptations in the NAc, contributing to deficits in extinction learning and memory consolidation, ultimately enhancing reinstatement of cocaine-seeking behavior.

The mPFC and NAc are particularly vulnerable to both cocaine and HIV infection, though the specific mechanisms by which circuit dysfunction modulates cocaine-seeking behavior in PLWH remain unclear. Circuit manipulation techniques, such as chemogenetics, can help reveal projection-specific approaches to reverse cocaine-induced dysfunction in the context of HIV infection. For example, chemogenetic activation of the ventromedial PFC-to-NAshell pathway has been shown to reduce cue-induced reinstatement of cocaine-seeking, suggesting that activating these circuits could be a potential strategy to mitigate relapse triggered by cocaine-associated cues (Augur et al., 2016). We found chemogenetic activation of the mPFC-NAshell pathway reduced reinstatement in EcoHIV-infected mice, suggesting that EcoHIV infection did not render reinstatement insensitive to mPFC-Nashell suppression of reinstatement. These findings suggest that promoting activation of mPFC-NAc circuitry is an effective strategy to mitigate cocaine seeking in EcoHIV-infected mice. Thus, despite facilitated reinstatement and altered Nashell cFos expression, targeted modulation of this circuit still has potential for the development of medications to treat CUD in the context of comorbid HIV. However, these results do not rule out the possibility of other circuits involving the NAshell in impaired extinction learning and enhanced cocaine-seeking behavior. The unique effect on cocaine-induced cFos expression by EcoHIV infection suggests that the observed effects may not be exclusive to the mPFC-NAshell pathway but could involve alternative circuits that were not investigated in this study.

A limitation of this study is a lack of precisely controlled AAV microinjection and DREADD expression across mPFC subregions (e.g., IL or PrL). Future study is necessary to dissect the precise circuit-level contributions to observed suppressing EcoHIV-induced reinstatement. This could be addressed using strategies such as retrograde targeting of the NAshell combined with optogenetic stimulation of the IL subregion. Overall, findings from the current study highlight the importance of incorporating circuit-level mechanisms into preclinical SUD research in PLWH which could accelerate the identification of effective targets for SUD prevention and treatment.

One additional consideration is the interpretation of the finding that EcoHIV-infected mice exhibited enhanced cFos induction in the NAshell in response to acute cocaine – suggesting elevated Nashell activity -, while activating glutamatergic inputs to the Nashell reduced reinstatement to cocaine seeking. One possibility is that selective activation of mPFC inputs to the NAshell is required for the modulation of reinstatement. It is also possible that cocaine-driven increases in NAshell activity could reflect alterations in dopaminergic activity that could compete with mPFC inputs (Aragona et al., 2008). Further, cFos expression captures a putative measure of activity across a broad temporal scale, and thus may not capture behaviorally-locked alterations in neural activity. Thus, the observed cFos expression suggests alterations in these substrates following EcoHIV infection and cocaine exposure, it likely does not represent the functional outcomes of neuronal activity in relation to cocaine-seeking behaviors. Further studies are needed to clarify EcoHIV-associated cell-type and circuit specific alterations in response to cocaine that underlying enhanced reinstatement to cocaine seeking.

Unexpectedly, EcoHIV infection did not affect yohimbine-induced reinstatement of cocaine-seeking. Yohimbine, an α2A-adrenergic receptor antagonist, functions by blocking the inhibitory effect of the α2A receptor on norepinephrine release, thereby increasing the stress- and anxiety-like responses (Charney et al., 1983). Yohimbine has been reported to induce cocaine craving and relapse (Mantsch et al., 2010; Feltenstein et al., 2011), and this phenomenon has been blocked by the α2A receptor agonist (Lee et al., 2004), implicating the α2A receptor in cocaine reinstatement. The studies examining HIV effects on α2A receptor are very limited. However, noradrenergic signaling in response to stress stimuli is impaired by HIV-associated immunodeficiency (Søndergaard et al., 2000). In addition, as HIV infection increased the dopaminergic neurotransmission in the CNS (Wang et al., 2004; Scheller et al., 2010), and dopamine (DA) is the precursor of norepinephrine and epinephrine, it is possible HIV-associated neuroimmune responses have indirect impacts on the noradrenergic system that contribute to stress activation. As no effect was observed on yohimbine-induced reinstatement of cocaine seeking, it is possible that EcoHIV did not affect α2A receptor at this experimental timepoint.

## Conclusion

We demonstrated that EcoHIV infection induced increased cocaine-primed reinstatement, which was associated differential activation of the NAshell. Chemogenetic activation of mPFC-NAshell projection Gq signaling was able to suppress cocaine-primed reinstatement of cocaine seeking in EcoHIV-infected mice as well as controls, which suggests that similar circuits may regulate drug seeking in EcoHIV infection. These findings provide valuable insights into potential strategies for managing cocaine relapse among people living with HIV.

## Acknowledgements

We would like to thank Dr. David Volsky for providing the EcoHIV-NDK plasmid. The authors would also like to thank Bryan Roth for providing the pAAV-hSyn-hM3D(Gq)-mCherry and Karl Deisseroth for the control virus pAAV-hSyn-mCherry. This research was supported by NIH award DP2DA051907 (JMB), R21DA056309 (JGJ), R01AG081929 (JGJ), F32DA060768 (MDN), and pilot awards from The Comprehensive Neuro-AIDS Center Grant P30MH092177-9 (JMB and MDN).

## Author contributions

Qiaowei Xie: Writing – review & editing, Writing – original draft, Methodology, Investigation, Formal analysis, Conceptualization. Rohan Dasari: Investigation. Mark D. Namba: Writing – review & editing, Writing – original draft. Lauren A. Buck: Writing – review & editing, Investigation. Christine M. Side: Investigation. Samuel L. Goldberg: Investigation. Kyewon Park: Methodology, Investigation. Joshua G. Jackson: Writing – review & editing, Methodology, Investigation. Jacqueline M. Barker: Writing – review & editing, Writing

– original draft, Project administration, Funding acquisition, Conceptualization.

